# Dynamics of global gene expression and chromatin accessibility of the peripheral nervous system in animal models of persistent pain

**DOI:** 10.1101/2021.01.27.427793

**Authors:** Kimberly E. Stephens, Weiqiang Zhou, Zachary Renfro, Zhicheng Ji, Hongkai Ji, Yun Guan, Sean D. Taverna

**Affiliations:** Department of Pediatrics, University of Arkansas for Medical Sciences, Little Rock, Arkansas; Arkansas Children’s Research Institute, Little Rock, Arkansas; Department of Biostatistics, School of Public Health, Johns Hopkins University, Baltimore, Maryland; Department of Biostatistics and Bioinformatics, Duke University, Durham, North Carolina; Department of Anesthesia and Critical Care Medicine, School of Medicine, Johns Hopkins University, Baltimore, Maryland; Department of Neurological Surgery, School of Medicine, Johns Hopkins University, Baltimore, Maryland; Department of Pharmacology and Molecular Sciences, School of Medicine, Johns Hopkins University, Baltimore, Maryland; Center for Epigenetics, Johns Hopkins University, Baltimore, Maryland

## Abstract

Efforts to understand genetic variability involved in an individual’s susceptibility to chronic pain support a role for upstream regulation by epigenetic mechanisms. To examine the transcriptomic and epigenetic basis of chronic pain that resides in the peripheral nervous system, we used RNA-seq and ATAC-seq of the rat dorsal root ganglion (DRG) to identify novel molecular pathways associated with pain hypersensitivity in two well-studied persistent pain models induced by Chronic Constriction Injury (CCI) of the sciatic nerve and intra-plantar injection of Complete Freund’s Adjuvant (CFA) in rats. Our RNA-seq studies identify a variety of biological process related to synapse organization, membrane potential, transmembrane transport, and ion binding. Interestingly, genes that encode transcriptional regulators were disproportionately downregulated in both models. Our ATAC-seq data provide a comprehensive map of chromatin accessibility changes in the DRG. A total of 1123 regions showed changes in chromatin accessibility in one or both models when compared to the naïve and 31 shared differentially accessible regions (DAR)s. Functional annotation of the DARs identified disparate molecular functions enriched for each pain model which suggests that chromatin structure may be altered differently following sciatic nerve injury and hind paw inflammation. Motif analysis identified 17 DNA sequences known to bind transcription factors in the CCI DARs and 33 in the CFA DARs. Two motifs were significantly enriched in both models. Our improved understanding of the changes in chromatin accessibility that occur in chronic pain states may identify regulatory genomic elements that play essential roles in modulating gene expression in the DRG.

**Summary:** Shared transcriptomic and epigenetic changes in two animal models improves our understanding of how chromatin structural changes alter DRG gene expression under persistent pain conditions.

## Introduction

Despite intense research efforts to develop novel analgesic classes, few novel molecular targets have been identified [15]. While non-steroidal anti-inflammatory drugs and opioids continue to be the most effective drugs commonly prescribed for the treatment of persistent pain, they have been associated with significant adverse effects. Therefore, increased attention is now focused on genetic and epigenetic mechanisms as an avenue to identify new druggable targets [22,24].

Current evidence supports the association between changes in gene expression and the transition from acute to chronic pain states in a number of preclinical and clinical models [17,33]. However, as these models are developed using nerve injury, administration of chemical agents, or evoking a significant inflammatory response, difficulties arise when disentangling gene expression profiles due to the effects of pain behaviors versus the initiating insults. Another factor that complicates the interpretation of transcriptional changes across different pain states is that existing microarray datasets from various preclinical models of persistent pain suffer from poor accuracy for genes expressed in low abundance and at lower coverage. Since these limitations can largely be overcome with newer RNA-seq approaches, the use of RNA-seq to identify shared changes in gene expression in disparate preclinical models is a promising strategy to identify pain-specific gene expression changes.

Chromatin structure is a well-known regulator of gene transcription across eukaryotic organisms [19,37]. However, chromatin structure in the peripheral nervous system (PNS) remains poorly understood. Intriguingly, recent work has indicated the involvement of epigenomic changes in the regulation of gene expression the dorsal root ganglion (DRG), which contains the soma of peripheral sensory neurons, in a preclinical model of neuropathic pain [26]. Long-term changes to gene expression patterns likely depend upon modifications to chromatin structure in post-mitotic neurons as well [36]. Thus, changes in chromatin structure at DNA regulatory regions in DRG neurons likely foster long-term changes in membrane potential and excitability, and thus, promote maintenance of persistent pain states. Histone acetylation and DNA methylation have been identified in persistent pain models, and our improved understanding of these and other epigenetic mechanisms through which aberrant gene expression could occur in the PNS unveil early molecular events that underlie the maintenance of chronic pain states and inform novel analgesic treatments.

Here, we used both RNA sequencing and the Assay for Transposase-Accessible Chromatin by sequencing (ATAC-seq) to identify patterns of genome-wide chromatin accessibility and gene transcription in the lumbar DRGs employing two widely used rodent models of neuropathic and inflammatory pain. The differences in chromatin accessibility observed between naïve and pain states in each model allowed us to identify regulatory DNA sequences and putative transcription factors that may drive the changes in gene expression associated with hyperalgesic states. Our integrative approach allowed us a greater understanding of the transcriptional and epigenetic basis of chronic pain in the DRG and identified novel biological processes and regulatory intermediates that may lead to the long-term transcriptional changes associated with persistent pain in the PNS.

## Methods

### Animals

Adult male Sprague-Dawley rats (12 weeks old; Harlan Bioproducts for Science, Indianapolis, IN) were allowed to acclimate for a minimum of 48 hours prior to any experimental procedures. Animals had access *ad libitum* to food and water. All procedures were reviewed and approved by the Johns Hopkins Animal Care and Use Committee and were performed in accordance with the National Institutes of Health’s Guide for the Care and Use of Laboratory Animals.

### Establishment of pain models

#### CCI of sciatic nerve

CCI surgery to the sciatic nerve was performed as previously described [3]. Under 2-3% isoflurane, a small incision was made at the level of the mid-thigh. The sciatic nerve was exposed by blunt dissection through the biceps femoris. The nerve trunk proximal to the distal branching point was loosely ligated with four 4-O silk sutures placed approximately 0.5mm apart until the epineurium was slightly compressed and minor twitching of the relevant muscles was observed. The muscle layer was closed with 4-O silk suture and the wound closed with metal clips.

#### Intraplantar injection of CFA

CFA (Sigma-Aldrich, St. Louis, MO) was diluted 1:1 with sterile 0.9% saline to produce a 0.5mg/ml emulsion. Under 2-3% isoflurane, the plantar surface of each hind paw was cleaned and injected with 100µl of the 50% CFA emulsion using a 27-guage hypodermic needle.

### ATAC-seq library preparation

Immediately following dissection, the ipsilateral lumbar (L4-L6) DRGs from one rat were transferred directly to cold lysis buffer (0.32M sucrose, 5mM calcium chloride, 3mM magnesium acetate, 10mM Tris-hydrochloride, pH 8.0, 0.1% Triton X-100, 1mM dithiothreitol, 5mM sodium butyrate, 1mM phenylmethylsulfonyl fluoride). Nuclei were isolated through dounce homogenization of the tissue in lysis buffer followed by sucrose gradient ultracentrifugation (1.8M sucrose, 3mM magnesium acetate, 1mM dithiothreitol, 10mM Tris-hydrochloride, pH 8.0, 5mM sodium butyrate, 1mM phenylmethylsulfonyl fluoride) at 139,800 x g at 4°C for 1 hour to remove mitochondrial DNA. The nuclei were resuspended in 1X phosphate buffered saline and counted 3 times using a Neubauer chamber. Tagmentation by Tn5 was performed using reagents from the Nextera DNA Sample Preparation Kit (FC-121-1030, Illumina; San Diego, CA) as previously described [5]. Each 50ul reaction contained 50,000 nuclei, 25ul 2X Tagmentation Buffer, and 2.5ul TDE1 enzyme and incubated at 37°C for 30 minutes. Tagmented DNA was immediately purified using the Clean and Concentrate-5 Kit (Zymo, Irvine, CA) and eluted in 10ul elution buffer. Tagmented DNA fragments were amplified using Nextera Index adapters, PCR primer cocktail, NPM PCR master mix and 10 cycles of PCR. Each library was purified using Agencourt AMPure XP beads (Beckam Coulter, Atlanta, GA). The fragment distribution of each library was assessed the using the High Sensitivity DNA Kit on an Agilent 2100 Bioanalyzer (Agilent Technologies, Palo Alto, CA). Libraries were quantified prior to sequencing using the Qubit DNA High Sensitivity kit (ThermoScientific, Waltham, MA) and normalized to 2nM and pooled in equimolar concentrations. Libraries were sequenced using paired end, dual-index sequencing on an Illumina HiSeq 2500 (Illumina, San Diego, CA) which produced 50 base pair reads.

### Data processing

The paired-end reads were trimmed using Trimmomatic [4] to remove adaptors. The trimmed reads were then aligned to rat genome rn6 using Bowtie2 [18] with the following parameters -X2000 --no-mixed --no-discordant. Reads with mapping quality score less than 10 were removed using SAMtools [20] and duplicated reads were removed using the MarkDuplicates function in Picard (http://broadinstitute.github.io/picard/). Aligned reads were shifted 4 nucleotides upstream for the 5’ end and 5 nucleotides downstream for the 3’ end to remove potential artifacts of Tn5 transposase binding. Tn5 transposase insertion sites were identified by trimming each read to the 5’ end. Bedtools *slop* was used to extend the insertion site by 75bp upstream and downstream [28]. Reads for each sample were downsampled to 49 million insertion sites to account for differences in sequencing depth. ATAC-seq peaks for each sample were called on the down-sampled bed files using *callpeak* function of Model-based Analysis of ChIP-seq (MACS2) and the parameters –nomodel –extsize 150 -B –keepdup all – call-summits [39]. Bigwig files were also generated from the down-sampled bed files for visualization in Integrative Genomics Viewer [32]. To improve the confidence of accessible regions in the dataset, peaks were considered for downstream analysis if the region was called in at least 50% of all samples from that group. A consensus peakset was then determined by the overlap of these regions.

The number of reads that aligned to each peak were counted and differential accessibility at each peak between the CFA and Naïve group and the CCI and Naïve group were determined using the *limma* package (version 3.38.3) in R (version 3.5.1) [31]. A p-value < 0.001 was used to define differentially accessible peaks between naïve and each of the chronic pain models. The genomic feature and the nearest annotated gene were determined using the *annotatePeaks*.*pl* function with the Hypergeometric Optimization of Motif Enrichment (HOMER) algorithm (version 4.11.1) [11]. De novo sequence motif discovery was to identify over representation of transcription factor binding sites within differentially accessible regions (DAR)s using the *findMotifsGenome*.*pl* function within HOMER [11].

### RNA-sequencing and data processing

#### RNA isolation

Total RNA was extracted from pooled ipsilateral lumbar (L4-6) DRGs from one rat using the Qiagen RNeasy Mini Prep Kit (Qiagen, Valencia, CA) with on-column DNase digestion (Qiagen, Valencia, CA) according to manufacturer’s instructions. RNA concentration was measured using the Nanodrop ND-2000 Spectrophotometer (Thermo Fisher Scientific, Waltham, MA) and RNA integrity was assessed using RNA Nano Eukaryote chips in an Agilent 2100 Bioanalyzer (Agilent Technologies, Palo Alto, CA).

#### RNA-seq library construction and sequencing

Five hundred nanograms of total RNA per sample was used to construct sequencing libraries (n=1 rat/sample). Strand-specific RNA libraries were prepared using the NEBNext Ultra Directional RNA Library Prep Kit for Illumina (New England Biolabs, Ipswich, MA) after poly(A) selection by the NEBNext poly(A) mRNA Isolation Module (New England Biolabs, Ipswich, MA) according to manufacturer’s instructions. Samples were barcoded using the recommended NEBNext Multiplex Oligos (New England Biolabs). Size range and quality of libraries were verified on the Agilent 2100 Bioanalyzer (Agilent Technologies, Palo Alto, CA). RNA-seq libraries were quantified by qPCR using the KAPA library quantification kit (KAPA Biosystems, Wilmington, MA). Each library was normalized to 2nM and pooled in equimolar concentrations. Paired-end sequencing was performed in a single lane on an Illumina HiSeq2500 (Illumina, San Diego, CA). Libraries were sequenced to an average depth of 33.9 million reads per sample.

#### RNA-seq data analysis

Sequencing reads were aligned to annotated RefSeq genes in the rat reference genome (rn6) using HISAT2 [14], filtered to remove ribosomal RNA, and visualized using the Integrative Genomics Viewer [32]. A gene count matrix that contained raw transcript counts for each annotated gene were generated using the *featureCounts* function of the Subread package in R [21] against the Ensemble rn6 transcriptome. This count matrix was then filtered for low count genes so that only those genes with >0 reads across all samples were retained. We relied on the automatic and independent filtering used by DESeq2 to determine the most appropriate threshold for removing genes with low counts [23].

To identify genes that were differentially regulated following nerve injury, raw transcript counts were normalized, log_2_ transformed, and analyzed using the default procedures in DESeq2 [23]. All downstream analyses on RNA-seq data were performed on data obtained from DESeq2. Adjusted p-values were corrected using the Benjamini-Hochberg method. An adjusted p-value <0.05 and an absolute log_2_ fold change > 0.5 were used to define differentially expressed transcripts between naive and each of the chronic pain models. DESeq2 adjusts for multiple testing by implementing the procedures of Benjamini and Hochberg [23]. Genes with differential expression between groups were then included in gene ontology (GO) and pathway analysis to infer their functional roles and relationships. GO analysis for enriched GO biological processes in each set of differentially enriched genes (DEGs) identified by DESeq2 was performed using Metascape [40]. We previously validated our RNA-seq data using qPCR in biological replicates [33].

### Cell culture

HEK293 cells were purchased from the American Type Culture Collection (CRL-1573™; Rockville, MD). Cells were maintained in Dulbecco’s modified Eagle’s medium (DMEM, Sigma, St. Louis, MO) supplemented with 10% fetal calf serum and penicillin-streptomycin and incubated at 37°C in a humidified environment containing 5% CO_2_. Cells with low passage numbers (i.e., <20) were used for all experiments.

### Cloning

Luciferase reporter constructs were constructed by cloning a candidate enhancer region into the pGL3 promoter vector (Promega; Madison, WI). Each region was inserted using standard restriction enzyme-based cloning techniques. The regions were obtained by PCR of rat genomic DNA. The 5’ end of the primers were modified to contain BglII (Forward primer) and MluI (reverse primer) restriction sites (Supplemental table 1). PCR was performed using the Pfu Turbo polymerase (Agilent Technologies; Palo Alto, CA) and touchdown thermocycling. The PCR products were digested and ligated into the BglII (AGATCT) and MluI (ACGCGT) restriction enzyme sites of the pGL3-Promoter luciferase vector (Promega; Madison, WI). The ligated products were transformed into chemically competent DH5α cells using ampicillin (100mcg/ml) to select for the recombinant plasmid-positive colonies. All constructs were verified by restriction enzyme digest and Sanger sequencing.

### Transfection and luciferase assays

HEK293 cells were seeded at 20,000 cells/well in 48 well plates in 250ul of complete media and grown to 60-70% confluence. Cells were then transfected with each reporter construct (450ng) and the 50ng pGL4.74 Renilla luciferase expression vector (Promega; Madison, WI) using ViaFect Transfection Reagent (Promega; Madison. WI) in 25ul Opti-MEM (ThermoScientific, Waltham, MA) with a 4:1 ratio in 250µl complete medium. The transfection efficiency of HEK293 cells was evaluated by transfecting cells with EGFP-N1 (Clontech; Mountain View, CA) in parallel reactions. 48 hours post transfection, Firefly and Renilla luciferase activities were measured using the Dual-Glo Luciferase reporter assay system (Promega). Firefly luciferase activities were normalized to the Renilla luciferase activity and expressed as the relative fold difference of the empty pGL3 promoter vector. Each luciferase construct was tested in quadruplicate.

### Data availability

Raw and processed sequencing data for all ATAC-seq data and the RNA-seq data for the CFA samples have been deposited in the NCBI GEO database under accession #GSEXXXX. The RNA-seq data files for the naïve and CCI groups were previously published [33] and are available under accession #GEO100122.

## Results

### 1. Differential gene expression changes in animal models of chronic pain

To determine how gene expression is altered in the lumbar DRG following the establishment of two widely used rat models of persistent pain, we compared RNA-seq data obtained 14 days following nerve injury (i.e., CCI model) to naïve rats and 48 hours following hind paw inflammation (i.e., CFA) to naïve rats (Figure 1A). Principal component analysis (PCA) shows clear segregation of the transcriptomes from the CCI and CFA pain models and naïve rats (Figure 1B). The first two principal components accounted for a total of 91%.

**Figure 1.**
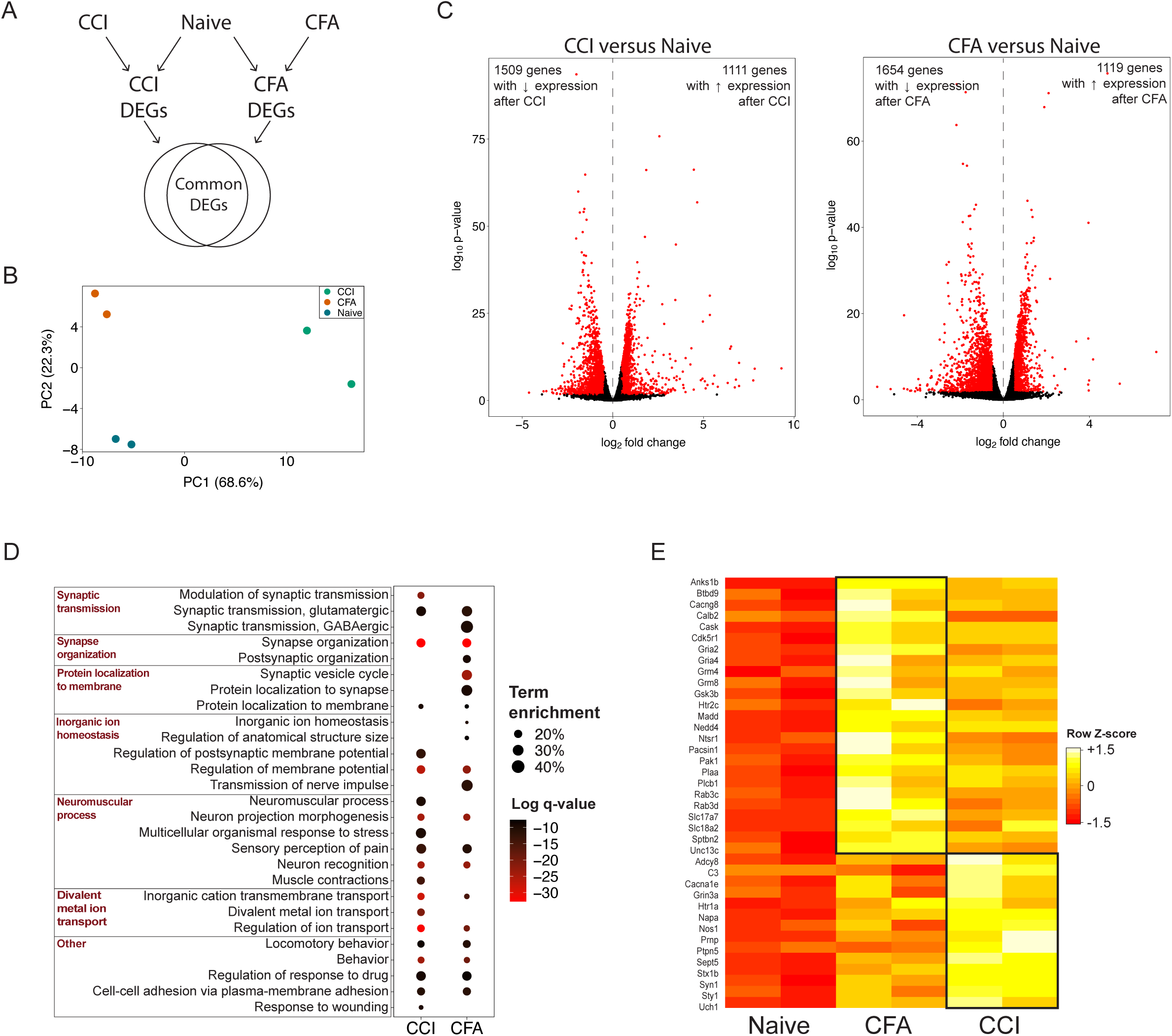
Changes in gene expression in the rat DRG in 2 models of chronic pain. A) Schematic of experimental setup. B) PCA plot of RNA-seq samples. C) Volcano plots of differential gene expression between CCI and naïve groups (left) and CFA and Naïve groups (right). Points highlighted in red indicated genes with differential expression as defined by an adjusted p-value < 0.05 and a log_2_fold change > |0.5|. D) Dot plot comparing the significant biological processes identified from the lists of upregulated genes differential expressed in each pain group (i.e., CCI compared to Naïve control, CFA compared to Naïve control). The log q-value for each term is indicated by color of each circle and the term enrichment (i.e., number of DEGs/total number of genes identified for that biological process) is indicated by size of the circle. E) Heatmap indicating CCI-and CFA-specific genes with roles in pre-synaptic plasticity that were significantly upregulated in each pain model compared to naïve control. CCI = Chronic Constriction Injury; CFA = Complete Freund’s Adjuvant; DEG = Differentially Expressed Gene.

Compared to naïve rats, we identified 2620 (17.8%) DEGs in the DRG following CCI. Of these 2620 DEGs, 1111 (42.4%) genes were upregulated following CCI as compared to naïve and 1509 (57.6%) genes were downregulated (Figure 1C). Gene ontology analysis of the 1111 upregulated genes indicated that neuronal-activity related biological processes including synapse organization, regulation of ion transport, modulation of chemical synaptic transmission and sensory perception of pain were among the biological processes that were statistically enriched (Figure 1D). Following CFA, we identified 2773 (18.8%) genes that were differentially expressed in the DRG when compared to naïve rats. Of these 2773 DEGs, 1119 (40.4%) genes were upregulated and 1654 (59.6%) genes were downregulated (Figure 1C). As expected, gene ontology analysis of the 1119 upregulated genes revealed neuronal- and pain-related biological processes including synaptic vesicle cycle, regulation of membrane potential, and transmission of nerve impulse were among the biological processes that were statistically enriched (Figure 1D). CCI- and CFA-specific changes in gene transcription that are important in pre-synaptic activity are shown in Figure 1E.

### 2. Shared differential expression of genes in CCI and CFA models

A total of 1960 genes were differentially expressed in both pain models with 752 genes upregulated and 1198 genes down regulated (Figure 2A; Supplemental table 2). GO analysis of the 752 upregulated genes showed significant enrichment among a variety of biological process related to synapse organization and membrane potential (Figure 2B top). GO analysis of the 1198 downregulated transcripts show significant enrichment among biological processes involved in transmembrane transport and ion binding (Figure 2B bottom).

**Figure 2.**
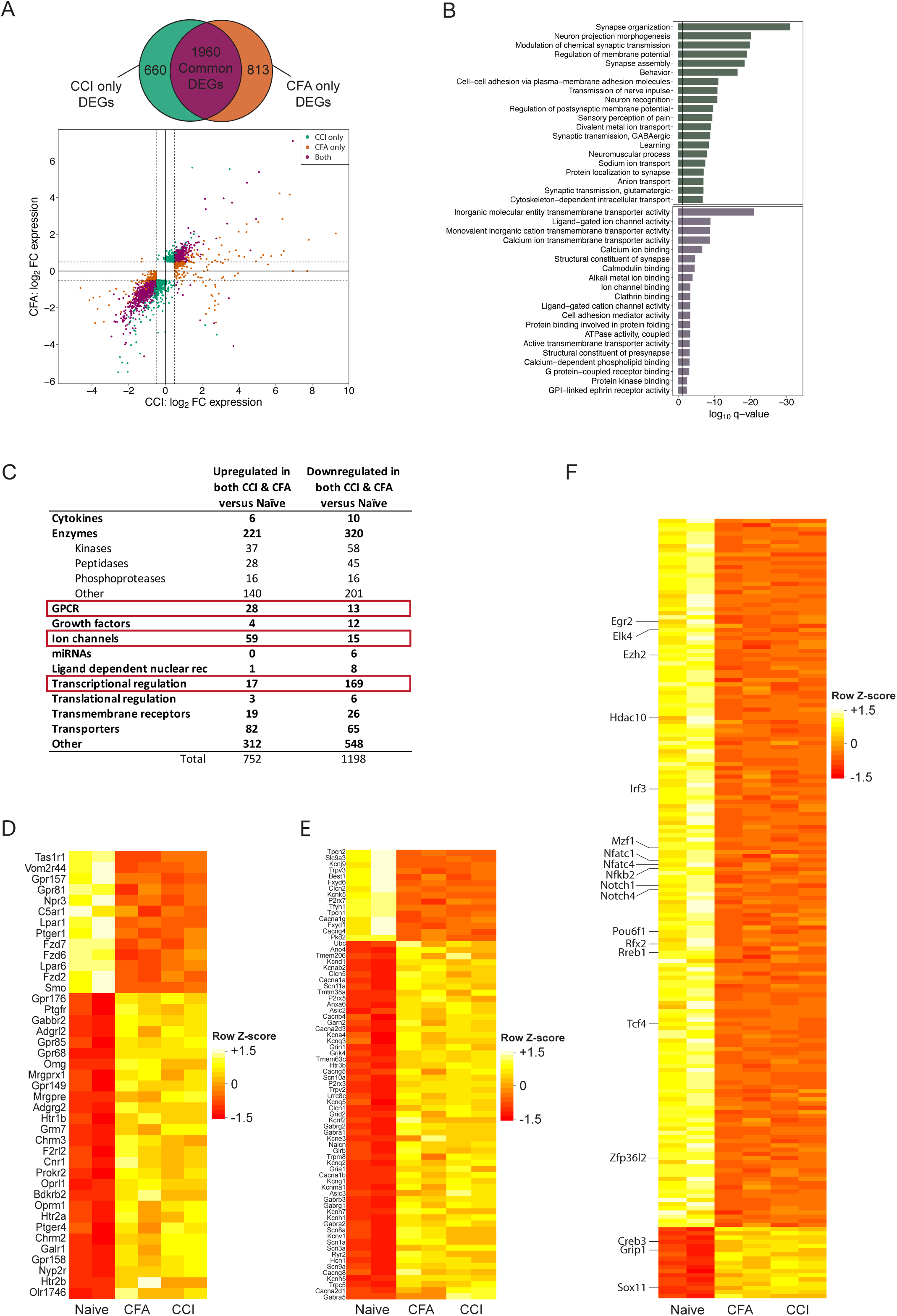
Shared DEGs following CCI and CFA in the rat DRG. A) Venn diagram shows the number of genes that are differentially expressed after CCI versus Naïve (green), CFA versus Naïve (orange), or in both models (purple). Log_2_FC expression of DEGs after CCI (x-axis) and after CFA injection (y-axis). Threshold of |log_2_FC| > 0.5 (dashed lines) with an FDR < 0.05 designates DEGs after CCI (green), CFA injection (orange), or DEGs found in both models (purple). B) Bar plot showing the top 20 GO biological processes significantly enriched in DEGs upregulated (top; green bars) and downregulated (bottom; purple bars) in both the CCI and CFA pain models. C) Table which shows the number of DEGs whose gene product is a member of each type of protein. For all common DEGs, the type of each gene product was identified for all DEGs upregulated and downregulated. A greater number of gene encoding GCPRs and Ion channels are upregulated in both models than the number downregulated. The data for these genes are provided in (D) and (E). A lower number of genes encoding transcriptional regulators are upregulated in both pain models than the number downregulated. The data for these genes are provided in (F). D-F) Heatmaps showing the DEGs that are in common following both CCI and CFA models of chronic pain for genes that encode GCPRs (D), ion channels (E), and transcriptional regulators (F). Colors in each row represent normalized gene counts for a single gene across the 2 biological replicates for each of the naïve, CFA and CCI groups. CCI = Chronic Constriction Injury; CFA = Complete Freund’s Adjuvant; DEG = Differentially Expressed Gene; FC = fold change; FDR = False Discovery Rate; GO = Gene Ontology

The numbers of upregulated genes that encoded GPCRs (Figure 2C,D) and ion channels (Figure 2C,E) were markedly higher than those of downregulated genes. Among the 28 GPCR genes upregulated following CCI and CFA, several are known to be involved in abnormal synaptic transmission of action potentials and altered pain thresholds (e.g., Gal1r, Gabbr2, Grm7, Gpr158, Chrm2, Chrm3, Cnr1, Oprl1, Oprm1, Mrgpre) as well as clinical pain conditions such as neuralgia (e.g., Htr2a, Htr2b, Ptger4, Mrgprx1) (Figure 2D). Conversely, of the genes identified as transcriptional regulators, the number downregulated was 10-fold higher than the number upregulated (Figure 2C,F; Supplemental table 3).

Of note, 10 genes (i.e., Myot, Ca3, Tnnc2, Ankrd23, Eno3, Casq1, Tpm2, Pygm, RT1-Da, Des) were upregulated after CCI and downregulated after CFA. The majority of these genes are involved in the function and maintenance of skeletal muscle and would be expected to be upregulated following our surgical dissection of the *biceps femoris*.

### 3. ATAC-seq provides a high-resolution chromatin accessibility profile of the rat DRG

Given our findings of gene expression changes in two persistent pain with different eitiologies, we hypothesized that these transcriptional changes were a result of dynamic chromatin occupancy in DRG cells which change the accessibility to cis-regulatory elements for transcriptional machinery. Therefore, we performed ATAC-seq of rat DRGs to determine the genome-wide dynamics of chromatin accessibility of the DRG in naïve rats and in the two pain models. Ninety to 94% of paired-end reads mapped to the rat reference genome and provided an average mapping depth of 17.8 million reads per sample. This represents an average of 35.6 million insertion sites per sample.

We identified a total of 97,485 peaks across all groups (i.e., Naïve, CCI, CFA). Between 53% and 69% of the peaks called in each group were found in only one sample in each group (Supplemental figure 1). Therefore, to increase our confidence that we were identifying true regions of open accessibility, we retained 56,810 accessible regions that were identified at least half of the samples in each group for use in all downstream analysis. With 33,628 regions (57.4% of consensus regions), we found that the CCI group had the smallest number of accessible sites. The CFA group had 46,238 (81.4%) accessible regions and the Naïve group had 45,399 (79.9%) accessible regions.

To determine the reproducibility among replicates, we performed principal component analysis and calculated Person correlation coefficients. We found that biological replicates were highly correlated with other samples from the same treatment group and less correlated with samples from either of the other 2 treatment groups. By PCA, samples show separation by group (Figure 3A). As expected, almost half (48.8%) of these regions were present in all three groups (Figure 3B). The distance between chromatin accessible regions and the nearest gene TSS suggests that these regions are concentrated in cis-regulatory regions (Figure 3C). Indeed, 54.7%, 28%, and 11.6% of all consensus accessible regions were located in intergenic, intronic, or promoter regions, respectively.

**Figure 3.**
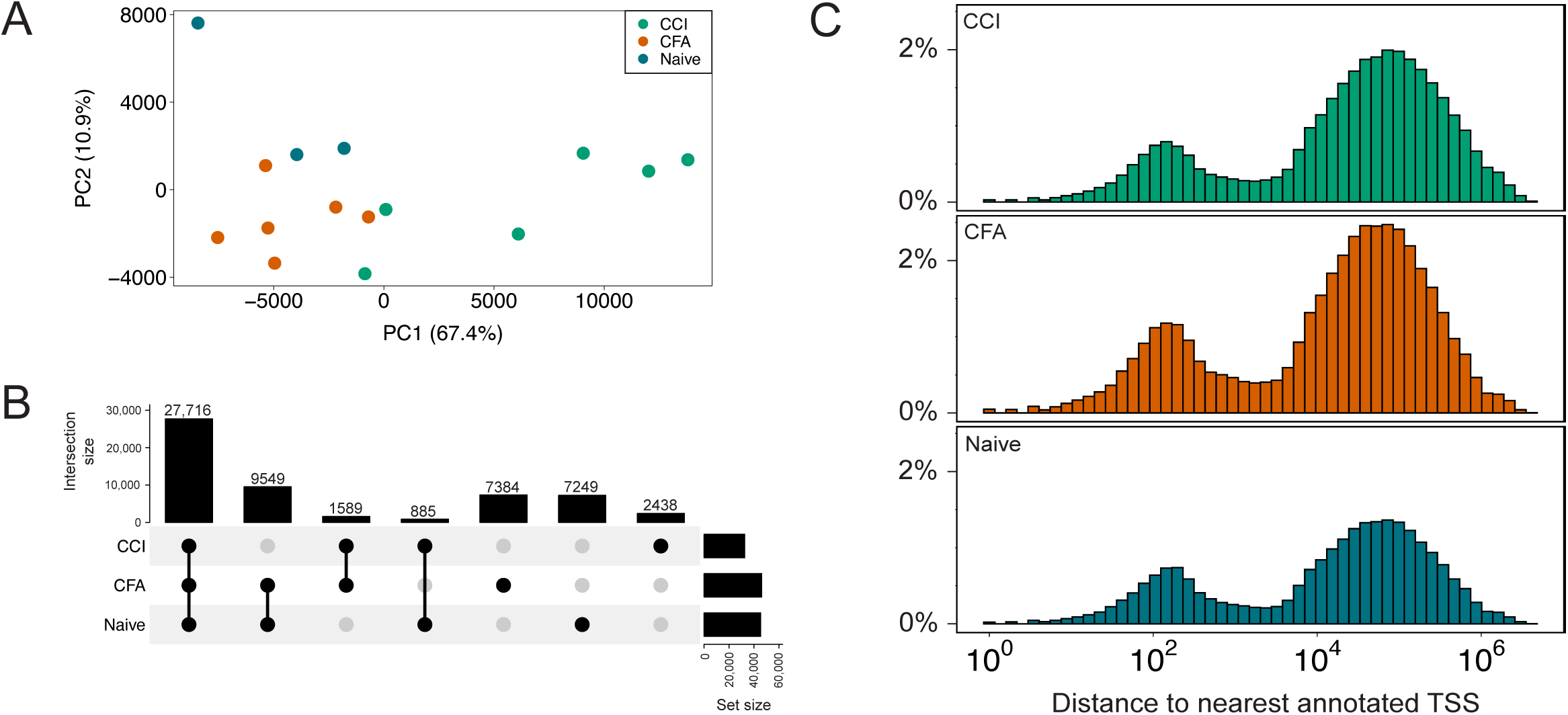
Chromatin accessibility in the rat DRG. A) Principal component analysis plot of all ATAC-seq biological replicates for each of the 3 groups. Percent of variance each component explained is included in axis labels. B) UpSet plot that shows the overlap of accessible regions among the 3 groups. C) Distance of accessible regions from nearest gene. The distribution of accessible regions shows concentration in cis-regulatory regions (i.e., gene promoter, intergenic/enhancer) versus uniform/random distribution along the genome. ATAC-seq = Assay for Transposase-Accessible Chromatin with high-throughput sequencing; CCI = Chronic Constriction Injury; CFA = Complete Freund’s Adjuvant; TSS = Transcription Start Site

### 4. Changes in chromatin accessibility in the DRG after the establishment of persistent pain

We evaluated each of the 56,810 consensus regions for changes in chromatin accessibility in each of the two pain models (p-value < 0.001). A total of 1123 (2.0%) of the 56,810 consensus regions showed changes in accessibility in one or both pain models compared to naïve rats. We found 517 DARs in the DRG following CCI compared to Naïve with 426 regions having increased accessibility and 91 regions having decreased accessibility (Figure 4A,B). When comparing the CFA model to naïve, we found 637 DARs with 321 regions with increased accessibility and 316 regions with decreased accessibility following CFA (Figure 4D,E). Sixty-two percent of all gains and losses in chromatin accessibility after CCI or CFA injection occurred in intergenic regions while 21.3% and 9.8% were located in introns and gene promoters, respectively (Figure 4G). These changes in genomic distribution of chromatin accessibility after sciatic nerve injury (CCI) or hind paw inflammation (CFA) may facilitate differential gene transcription through chromatin-level regulation at cis-regulatory regions.

**Figure 4.**
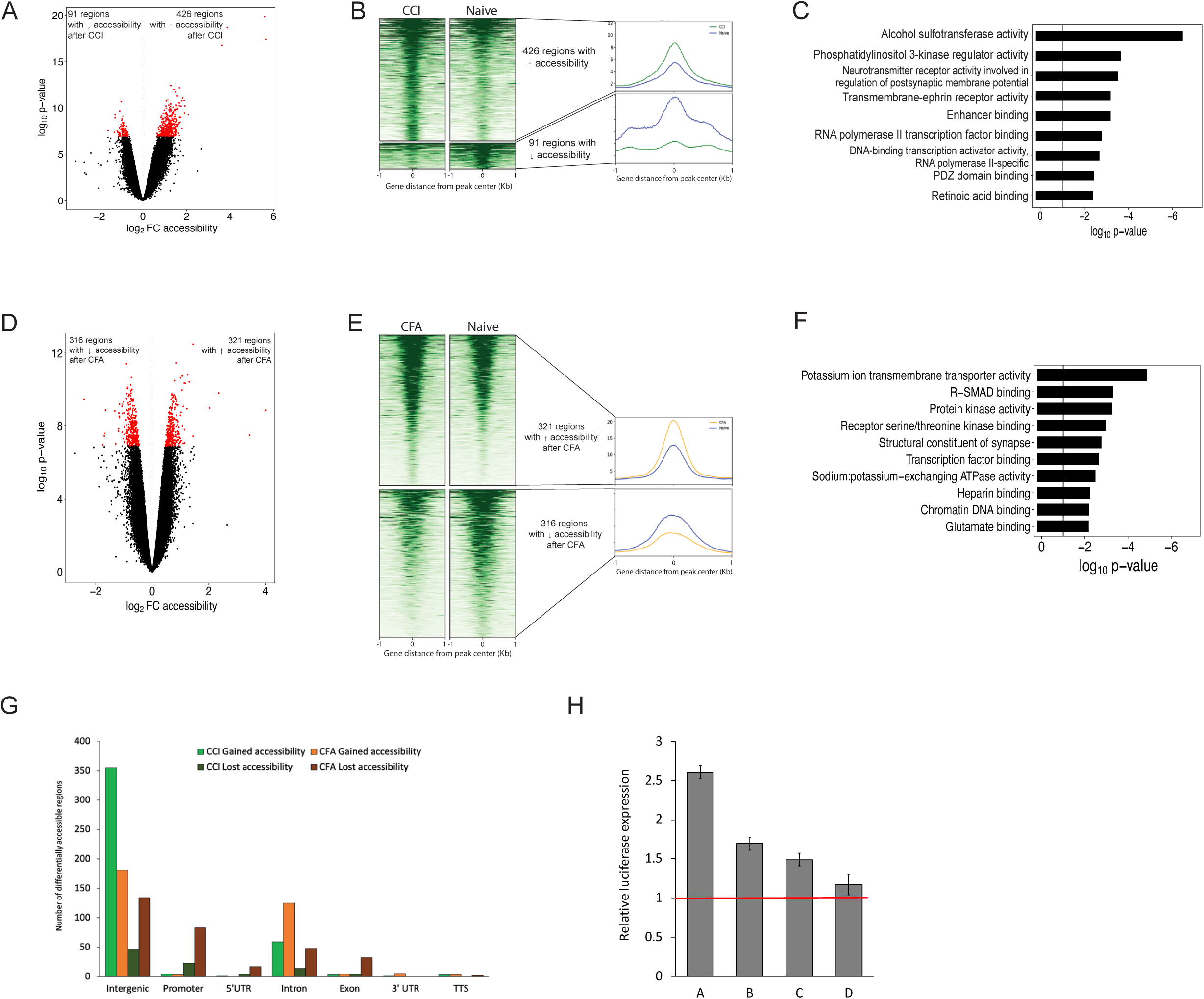
Chromatin accessibility in the rat DRG after CCI and CFA. Volcano plots of DARs after CCI versus naïve groups (A) and after CFA versus Naïve (D). Points highlighted in red indicate genes with differential accessibility as defined by a p-value < 0.001. Heatmap of read density for all DARs in CCI and Naïve (B) and CFA and Naïve (E) groups with increased accessibility (top panels) and decreased accessibility (bottom panels). Each row represents one accessible region and the regions are aligned at the center of each region. The color represents the intensity of chromatin accessibility. The average read density for each heatmap is shown to the right. C-F) Gene ontology analysis of the molecular functions identified in DARs after CCI versus Naïve (C) and CFA versus Naïve (F). G) Bar graph showing the proportion of DARs across genomic features after CCI or CFA. H) Activity of candidate DRG enhancers by luciferase reporter gene assays. Individual reporter plasmids were prepared as described and contained one candidate enhancer regions (A -D; Supplemental table 1). Luciferase activity was normalized to that of the Renilla reporter and expressed as mean fold relative activity of the empty reporter ± standard deviation. All constructs were tested in quadruplicate. CCI = Chronic Constriction Injury; CFA = Complete Freund’s Adjuvant; DAR = Differentially Accessible Region

Functional annotation of the DARs after CCI shows enrichment for molecular functions involved in phosphatidylinositol 3-kinase regulator activity, neurotransmitter receptor activity involved in the regulation of postsynaptic membrane potential and enhancer binding (Figure 4C). These functions converge on mechanisms with the potential to alter neuronal excitability and chromatin structure in DRG cells to produce pain hypersensitivity. Following CFA injection, enrichment in potassium ion transmembrane transport, receptor serine/threonine kinase binding, and structural constituent of synapses are among the molecular functions found in DARs (Figure 4F). The different molecular functions identified through GO analysis suggests that chromatin structure may be altered to different effect following nerve injury and hind paw inflammation.

A total of 31 DARs were shared following CCI and CFA injection (Table 1). Ten DARs showed decreased accessibility after CCI and CFA compared to Naïve. Of these 10 DARs, 2 were associated with upregulation of the nearest annotated gene (i.e., Grik4, Agtpbp1) and 2 were associated with down regulation of their nearest gene (i.e., Wdr60, Dlg3). A total of 21 DARs showed increased accessibility after CCI and CFA compared with Naïve. Of these 21 DARs, 3 were associated with upregulation of the nearest gene (i.e., Rrmb, Arap2, Pcdh9) and 2 were associated with downregulation of the nearest gene (i.e., Kif13b, Lpar1). The ability of these regions to modulate gene transcription were validated by luciferase assay (Figure 4H).

To determine transcription factors that may have their binding sites within with regions of chromatin accessibility in each of the pain models, we performed motif analyses using HOMER on all of the DARs identified in each pain model (Figure 5A and Supplemental table 4 and 5). We found a total of 17 DNA sequences known to bind transcription factors in the CCI DARs and 33 in the CFA DARs. Two binding motifs were significantly enriched following both CCI and CFA (Figure 5B). CFA was associated with the induction of a wider remodeling of transcription factor binding sites in the DRG than CCI. These enriched sites found after CFA clustered into several important families (e.g., high-mobility group (HMG), basic leucine zipper (bZIP), Transcriptional enhancer factor (TEF)). CCI was associated with enriched sites clustered into the ETS-domain (ETS) and zinc finger (Zf) families. The changes in the availability of potential TF binding sites support the long-lasting effects of our pain models on chromatin structure.

**Figure 5.**
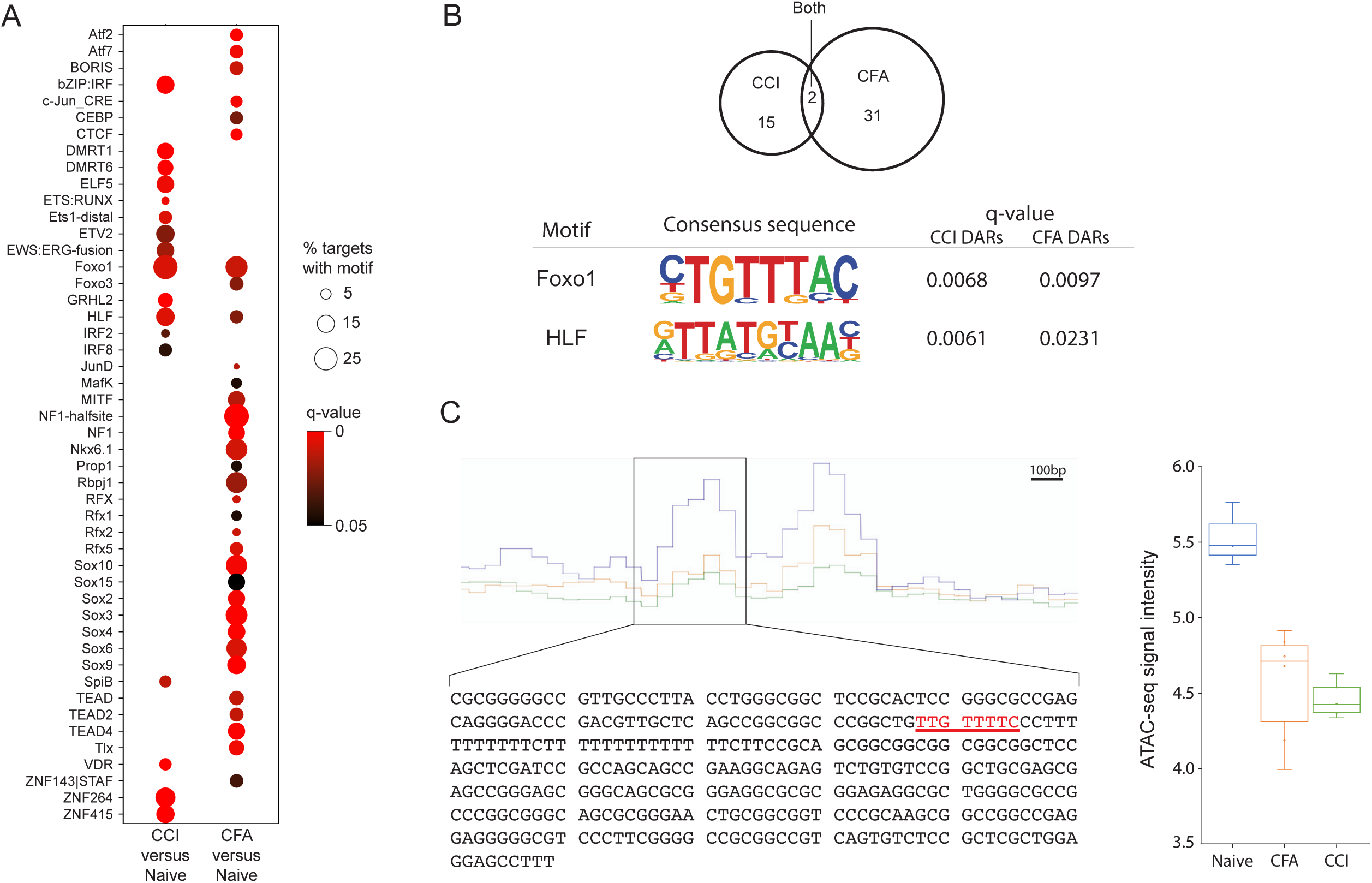
Motif analysis reveals shared DNA binding sequences. A) Dot plot of the significantly overrepresented motifs in DARs after CCI and CFA. The size of the circle represents the % of DARs that contain the motif and color indicates the q-value. B) Venn diagram that shows the overlap of the number of DNA binding motifs in the DARs between CCI and Naïve, and the CFA versus Naïve comparisons. Two motifs are common which are identified as Foxo1 and Hlf. The motif and statistics from HOMER for each of these motifs is provided. C) Chromatin accessibility at a putative regulatory region of the Grik4 gene. The average ATAC-seq signal of the downsampled, normalized bigwig files for each group as displayed from the Integrated Genomics Viewer (left). The region identified as being differentially accessible in both the CCI and CFA groups compared to the Naïve group is outlined (chr8:47,237,180-47,237,538). This 358bp sequence is provided with the putative Foxo1 binding site highlighted in red. A box plot of the normalized log_2_transformed ATAC-seq signal for each sample (naïve, n=3; CFA, n=6; CCI, n=5). CCI = Chronic Constriction Injury; CFA = Complete Freund’s Adjuvant; DAR = Differentially Accessible Region

## Discussion

In this study we used both RNA-seq and ATAC-seq on DRG tissues to improve our understanding of the transcriptomic and epigenetic mechanisms in the PNS that may underlie persistent pain. We also aimed to identify novel molecular pathways involved in the development of pain hypersensitivity in two well-studied rodent models of persistent pain with different etiologies. Prior research using preclinical pain models has primarily relied on microarrays to study gene expression changes in the DRG [17]. The decreasing cost of next-generation sequencing has resulted in adoption of RNA-seq as the current standard which has a greater dynamic range for gene expression detection, the ability to measure a larger number of gene transcripts, and can detect differences in sequence and isoforms. Studies that have used RNA-seq to look at DRG changes, have primarily examined peripheral nerve injury models [9,12,27,34] with few examining non-nerve injury models (e.g., diabetic neuropathy [2], ultraviolet-induced inflammation [6]) which makes it difficult to identify gene expression changes that are specific to pain process versus nerve injury or specific disease-related processes.

In this study, we performed RNA-seq using two widely used persistent pain models which are induced by sciatic nerve CCI and hind paw inflammation to identify pain-related gene expression changes in the DRG. Consistent with prior findings, both nerve injury and inflammatory pain were found to be associated with transcriptional changes of genes involved in cell signaling (i.e., GPCR function, ion channel expression, synaptic transmission). Further, our findings were consistent with previous transcriptome-wide screens that support the upregulation of Reg3b, Vgf, Ccl2, P2rx3, Crh, Scn11a, Drd2, Npy2r, Cacna2d1, and Neto1 in the DRG in various pain models [6,17,27,29]. In addition, our findings are consistent with prior work that found genes downregulated (e.g., Rlbp1, Gja1, Lgr5, Lpar1, Ttyh1) in DRG neurons after CCI [29]. Here, we confirm that these genes are also downregulated in an inflammatory pain model which suggests that these genes have broader roles in pain pathways outside of nerve injury induced neuropathic pain.

Our studies identify several novel genes with previously unknown functions in the development and maintenance of pain. For example, Plxna2 was upregulated in both CCI and CFA models. Plxna2 encodes the plexin-A2 receptor known to be expressed in hippocampal and cortical neurons [10,35] Upon binding by its ligands, semaphorin-3A or -6A, plexin-A2 triggers an intracellular signaling cascade which mediates axon repulsion and cell migration during nervous system development [25,30]. Further research is needed to determine how Plxna2 may participate in pronociceptive mechanisms.

Interestingly, we found that genes involved in epigenetic and transcriptional regulation were largely downregulated following both CCI or CFA injection. Consistent with prior evidence we found that Dnmt3a, Dnmt3b, Sirt2, Brd3, and Ehmt2 (which encodes G9a) were among the genes downregulated in both pain models although the magnitude of the log_2_ fold change did not meet our criteria to be significant in the present study [8,13,38]. Mounting evidence supports the roles for epigenetic mechanisms and our findings that a large proportion of transcriptional regulators are downregulated supports this idea. Genes that decrease the regulation of existing transcriptional programs promote the transcriptional changes which are now well established in preclinical and clinical models of persistent pain. These changes in gene transcription and transcriptional regulation may facilitate neuronal hyperexcitability-induced remodeling of chromatin structure and neuronal responsivity of cells in the DRG. Our findings are consistent with prior studies which provides indirect evidence of decreased transcriptional control in persistent pain states. HDAC inhibitors and HAT show analgesic effects in various pain models through non-specific changes in chromatin structure.

One of the limitations in the study of epigenetic mechanisms in the PNS is the small number of cells that comprise each DRG. Traditional chromatin accessibility assays require millions of cells. However, ATAC-seq can assess native chromatin accessibility with much less starting material, and therefore, can detect subtle changes in chromatin accessibility both in homogenous cell lines and in heterogenous mixtures of cell types [5].

Our study is the first to provide a comprehensive map of changes in chromatin accessibility in the PNS using both neuropathic pain and inflammatory pain models. Chromatin accessibility is necessary for transcription factor binding to cis-regulatory regions and subsequent changes in gene expression [16,37]. The DRG is a collection of several types of primary sensory neuronal cell bodies and satellite glia cells which acts as the initial point of modulation of action potentials from potentially noxious stimuli. Gene expression changes in the DRG control noxious input to second order neurons in the spinal cord. Therefore, understanding changes in chromatin accessibility that occur in chronic pain states would help to identify regulatory loci in the DRG that play a central role in changing gene expression there. We identified 1123 regions that showed dynamic chromatin accessibility after CCI and/or CFA. Within these regions we found 48 known DNA binding motifs for transcription factors. Importantly, we identified overrepresentation for the DNA binding motifs of two transcription factors in DARs after both CCI and CFA injection. For example, we found that in both the CCI and CFA models, decreased chromatin accessibility at a 358bp region ∼130Kb downstream of Grik4 is associated with increased Grik4 gene expression (Figure 5C). Grik4 encodes a subunit of a glutamatergic receptor that contributes to excitatory postsynaptic currents and is expressed in the central terminals of nociceptive neurons that synapse in laminae I-III [7]. Over expression of Grik4 in the mouse brain is associated with increased amplitude, greater frequency and quicker decay of spontaneous EPSCs in CA3 cells [1]. While the effects of increased Grik4 expression in DRG neurons has not been investigated, these alterations in synaptic transmission are consistent with the increased efficacy of synaptic transmission of nociceptive input of central sensitization. The 358bp DAR we identified upstream of Grik4 in the rat maps to chr11:120511578-120511909 on hg19 and is located within a previously identified regulatory region for Grik4 (i.e., GeneHancer #GH11J120511). Our findings suggest that this site is a transcriptional repressor binding site and that decreased accessibility inhibits this repression to increase Grik4 gene expression. Indeed, we found a potential Foxo1 binding site within this DAR (Figure 5C). Foxo1 is well established as a transcriptional repressor in neural tissues. Further research is needed to evaluate this region for regulatory potential and determine if Foxo1 binds to this region in the rat DRG and can modulate pain behaviors.

Our bulk ATAC-seq provides an average of chromatin accessibility at a specific genomic locus. Change in chromatin accessibility may be due to an increase in the number of cell types in which a specific region is accessible or an increase in accessibility in the existing population of cells [16]. As low-cell number methodologies mature, future research may identify the epigenetic mechanisms responsible for the changes in chromatin structure in specific cell types.

**Table 1.**
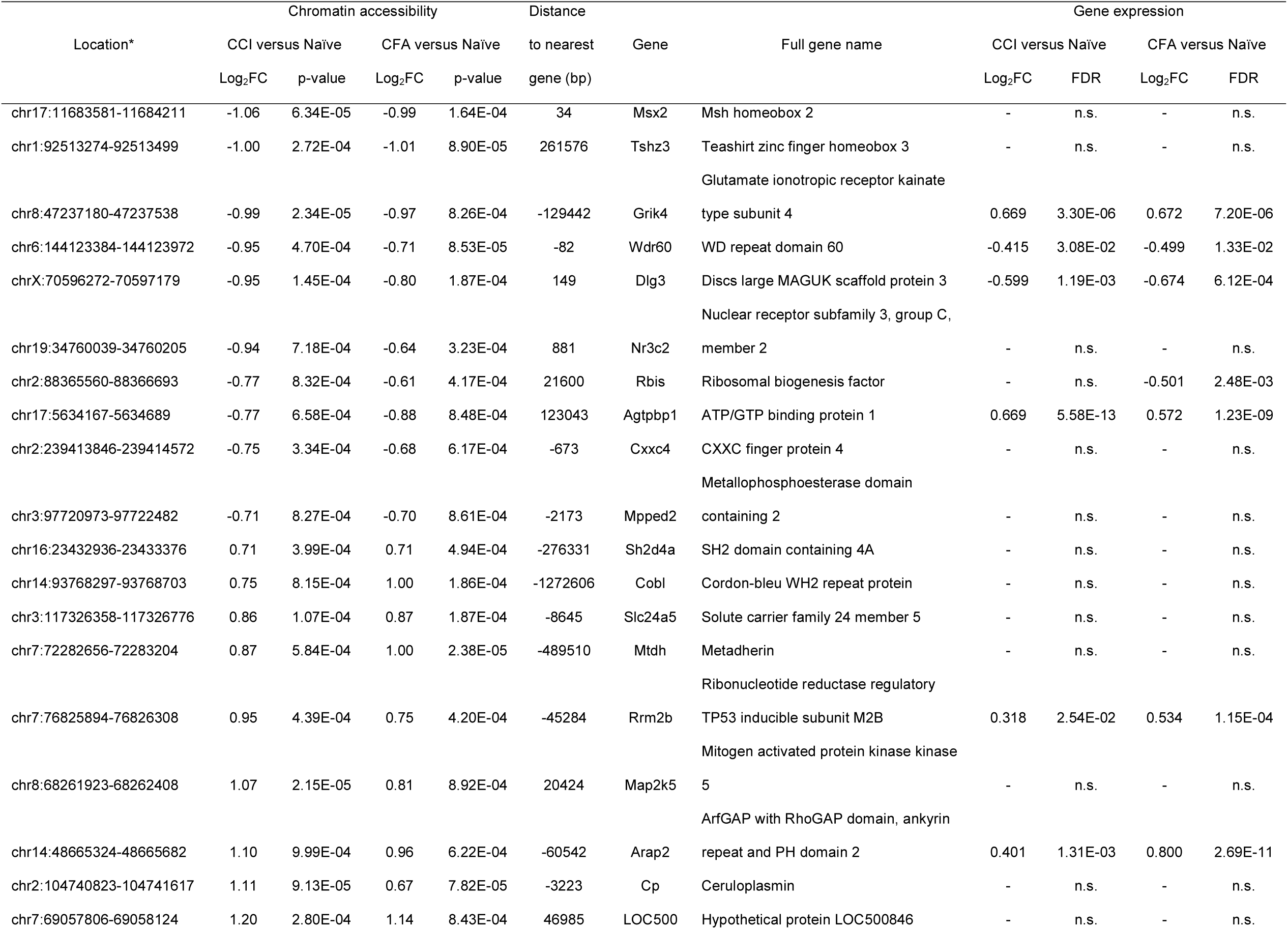
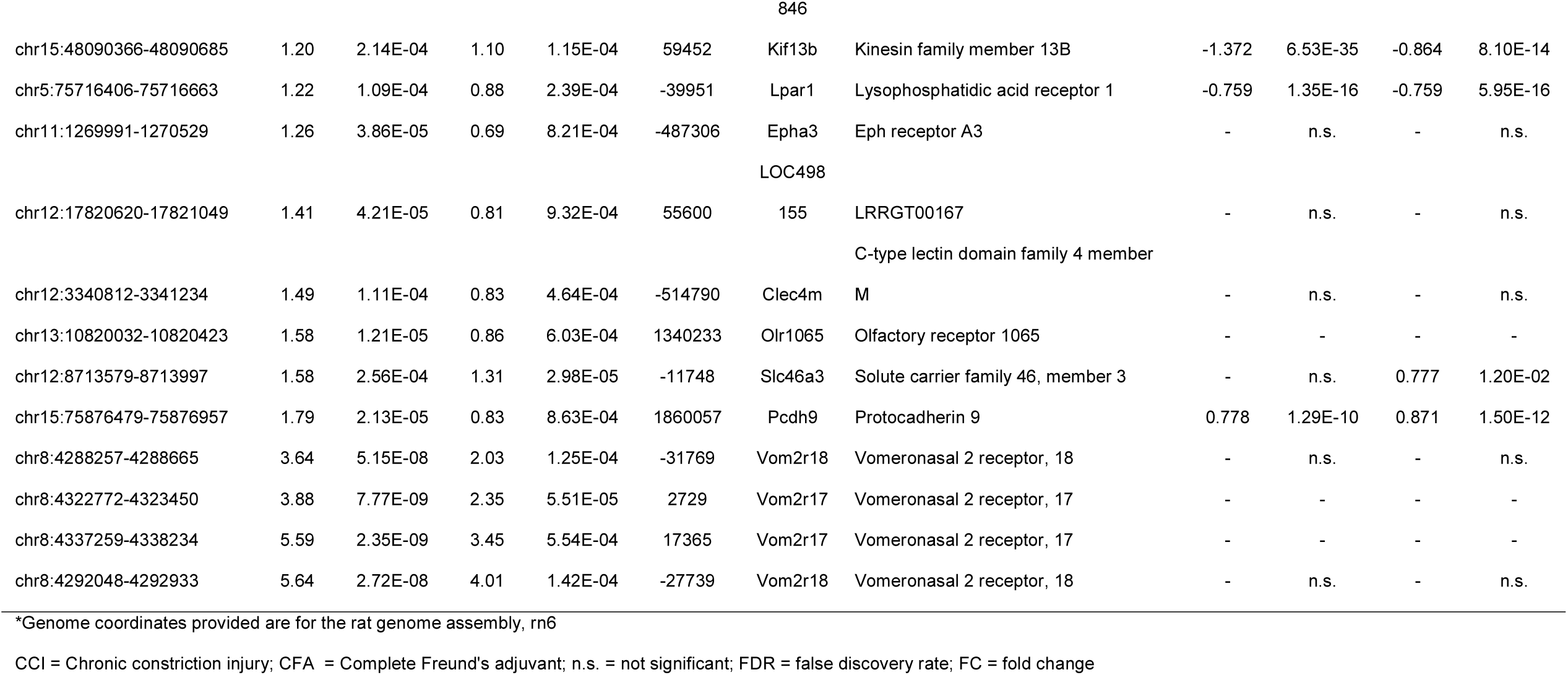
Differentially accessible regions in the dorsal root ganglion following either CCI or CF

## Acknowledgements

This study was supported by grants from National Institutes of Health (Bethesda, Maryland, USA) F32NR015728 (KES), KL2 TR003108 (KES), NS110598 (YG), NS117761 (YG), R01GM118760 (SDT), the Arkansas Children’s Research Institute (KES), the Arkansas Breast Cancer Research Program (KES) as well as a seed grant from the Johns Hopkins Blaustein Pain Research Fund (SDT). We thank Rakel Tryggvadóttir and Dr. Jaclyn Daniels for their technical assistance. All authors report no conflicts of interest.

---

**Supplemental figure 1. Consistency of peaks among replicates.** The graph shows the number of peaks identified in one or more samples. Due to the large number of accessible regions identified in only one sample, we included only regions that were identified in at least 50% of the samples within the study group.

## Notes

### Competing Interest Statement

The authors have declared no competing interest.

