## Supplementary figures and images for "Dynamics of global gene expression and chromatin accessibility of the peripheral nervous system in animal models of persistent pain"

### Supplemental figure 1

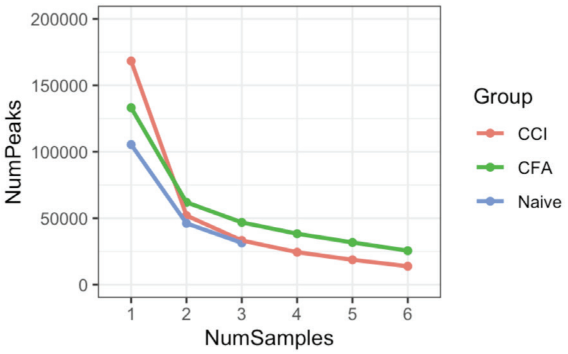
